# Active surveillance of cats and dogs from households with human COVID-19 cases reveals over one quarter of pets infected with SARS-CoV-2 in 2020-2021 in Texas, United States

**DOI:** 10.1101/2025.02.03.636361

**Authors:** Alex Pauvolid-Corrêa, Edward Davila, Lisa D. Auckland, Italo B. Zecca, Rachel E. Busselman, Wendy Tang, Christopher M. Roundy, Mary Lea Killian, Mia K. Torchett, Melinda Jenkins-Moore, Suelee Robbe Austerman, Kris Lantz, Katherine Mozingo, Rachel Tell, Ailam Lim, Yao Akpalu, Rebecca S. B. Fischer, Francisco C. Ferreira, Gabriel L. Hamer, Sarah A. Hamer

**Affiliations:** Department of Veterinary Integrative Biosciences, Texas A&M University, College Station, TX 77843, USA; Department of Entomology, Texas A&M University and AgriLife Research, College Station, TX 77843, USA; National Veterinary Services Laboratories, USDA APHIS VS, Ames, IA 50010, USA; Wisconsin Veterinary Diagnostic Laboratory, University of Wisconsin-Madison, Madison, WI 53706, USA; Brazos County Health Department, Bryan, TX 77803, USA; School of Public Health, Texas A&M University, College Station, TX 77843, USA

**Keywords:** SARS-CoV-2, One Health, risk factor, COVID-19, United States, Texas, cat, dog

## Abstract

Households where people have COVID-19 are high risk environments for companion animals that are susceptible to SARS-CoV-2. We sampled 579 pets from 281 households with one or more laboratory-confirmed person with COVID-19 in central Texas from June 2020 to May 2021. Nineteen out of 396 (4.8%) dogs and 21 out of 157 (13.4%) cats were positive for SARS-CoV-2 by RT-qPCR. Additionally, 95/382 (25%) dogs and 52/146 (36%) cats harbored SARS-CoV-2 neutralizing antibodies. Twenty-six companion animals of ten other species were negative. Overall, 164 (29%) pets were positive for SARS-CoV-2 by molecular and/or serological tests; a total of 110 (39%) out of 281 households had at least one animal with active or past SARS-CoV-2 infection. Cats were more likely to be infected by SARS-CoV-2 and had higher endpoint antibody titers than dogs. Through viral isolation from a subset of respiratory swabs, we documented 6 different lineages in dogs and cats, including the B.1.1 lineage in a cat one month prior to the first known human case in the country. We observed animal and human-pet interaction factors associated with higher risk of infection for dogs and cats, such as days after COVID-19 diagnosis and sharing food. Frequency of clinical signs of disease reported by owners of pets with active infections did not differ from uninfected ones, suggesting that not all reported signs are attributed to SARS-CoV-2 infection. Characterizing animal infections using active SARS-CoV-2 surveillance in pets at risk of infection may aid in One Health pandemic prevention, response, and management.

## Introduction

Severe acute respiratory syndrome coronavirus 2 (SARS-CoV-2), currently classified as *Betacoronavirus pandemicum,* is a zoonotic virus species that spread globally, and has generated thousands of variants [1,2]. While the main driver of COVID-19 transmission is human-to-human, cases of SARS-CoV-2 infection in animals resulting from spillback from infected people have been reported in multiple countries [3]. SARS-CoV-2 has a broad range of susceptible mammalian host species, and animal infections are presumed to result from close contact between infected people and animals [4].

Among all the wild, domestic, and captive animals that can be infected, pets living in households with active human infections represent are particularly at-risk because of their prolonged, close proximity and indoor relationships with humans [5,6]. Therefore, it is critical to understand factors associated with higher risk of human-to-pet transmission to protect the health of companion animals. Additionally, understanding the association between exposure to SARS-CoV-2 and the development clinical signs of disease in pets is important to inform owners and veterinarians about the consequences of infection.

In summer 2020, we initiated an active household surveillance study in east-central Texas in homes with people with COVID-19, documenting several infected cats and dogs, including animals with infectious virus in their respiratory tract [7]. We further showed that household pets are infected with the same SARS-CoV-2 strains that circulate in humans with the first report of the Alpha variant in companion animals in the United States [8]. Here, we build upon this framework with the complete results of our study that spanned over a year and sampled 579 pets with the objectives of characterizing the exposure and clinical presentations of dogs and cats throughout the initial phases of the pandemic, as well as the risk factors of animal infections.

## Materials and methods

### Animal recruiting and sampling

We sampled companion animals in Texas within the first 1.5 years of the pandemic, from June 2020 to May 2021. Sampled animals lived in the same household as a person with a laboratory-confirmed SARS-CoV-2 positive test, as previously described [7]. Households were identified by the Brazos County Health Department (BCHD) as part of a public health case investigation process. This project was approved by the Texas A&M University Institutional Animal Care and Use Committee and the Clinical Research Review Committee (2018-0460 CA). Companion animal owners completed a questionnaire by phone where they were asked for information regarding their animals including demographic information, animal care, and animal level of contact with the positive human. Additionally, owners were asked to report clinical signs of disease in their pets within two weeks of human diagnosis. Project personnel were deployed to the households for animal sampling, including collection of oral, nasal, rectal and body (fur) swabs collected in tubes containing viral transport media (VTM) and later stored at −80◦C. Oral and nasal swabs were initially combined into the same tube as a single respiratory sample, but were later collected separately. A subset of cats also had conjunctival swabs collected. Blood was collected after which serum was separated and stored at −80◦C. A subset of animals were sampled twice or more. Resampling was triggered by the detection of at least one positive animal in the household during the initial sampling event in all cases but in two households, and all pets in the house were sampled upon subsequent events.

### Molecular testing

Swab VTM samples were evaluated for the presence of SARS-CoV-2 viral RNA at TAMU and the Wisconsin Veterinary Diagnostic Laboratory (WVDL). Briefly, viral RNA was purified using commercial nucleic acid extraction kits (MagMAX CORE Nucleic Acid Purification Kit), as previously described [7]. All VTM samples were tested on separate specific quantitative reverse transcription polymerase chain reaction (RT-qPCR) targeting various genes of SARS-CoV-2, including the RNA-dependent RNA polymerase (RdRp), envelope (E), nucleocapsid region 1 (N1) and/or nucleocapsid region 2 (N2) [9–12]. From June to December 2020, all samples that tested positive by RT-qPCR were sent to the National Veterinary Services Laboratories (NVSL) of the U.S. Department of Agriculture (USDA) for confirmation with joint approval from the Texas Department of State Health Services and the Texas Animal Health Commission and in compliance with the World Organization for Animal Health (WOAH) reporting procedures [13]. Starting in January 2021, only samples that had Cycle thresholds (Ct) values less than 35 were sent to NVSL for confirmation.

### Viral isolation and plaque assay

Selected VTM samples that were positive by RT-qPCR for at least two genes in the same laboratory were subjected to virus isolation at TAMU and/or NVSL, as previously described [7,8]. Briefly, samples diluted from 1:2 to 1:3 in cell culture medium were inoculated in Vero (ATCC CCL-81) cells for viral adsorption followed by incubation at 37◦C from four to up to seven days in a total of two to three blind passages. Samples that presented cytopathic effect (CPE) in cell culture were confirmed positive for SARS-CoV-2 by RT-qPCR followed by Sanger nucleotide sequencing.

### Whole genome sequencing and phylogenetic analysis

At NVSL, libraries for whole-genome sequencing were generated from samples that were RT-qPCR positive for SARS-CoV-2, prioritizing samples with values Ct <30, using the Ion AmpliSeq Kit for Chef DL8 and Ion AmpliSeq SARS-CoV-2 Research Panel (Thermo Scientific, Waltham, MA, USA). Libraries were sequenced using an Ion 520 chip on the Ion S5 system using the Ion 510™ & Ion 520™ & Ion 530™ Kit. Sequences were assembled using IRMA v. 0.6.7 and visually verified using DNAStar SeqMan NGen v. 14.

### Virus neutralization

Serum samples were tested for specific neutralizing antibodies at NVSL by an in-house virus neutralization (VN) [13] and/or by a SARS-CoV-2 Surrogate Virus Neutralization Test (sVNT; GenScript, N.J., USA). For VN, two-fold serially diluted sera from 1:8 to 1:2,048 were challenged in duplicate with 100 TCID_50_/mL of SARS-CoV-2 (2019-nCoV/USA-WA1/2020; ATCC NR-52281) in Vero 76 cells (ATCC CRL-1587). The VN titer of a sample was recorded as the reciprocal of the highest serum dilution that provided full neutralization of the reference virus, as determined by visualization of CPE. Three cats with titers exceeding 1:512 did not have their endpoint titers established, in which case 512 was used for statistical analyses. For sVNT, the optical densities of the reactions of the test sera and the positive and negative controls supplied were read at 450 nm (OD450), and percentage inhibitions were calculated as follows: percent inhibition = (1 – sample O.D. value/negative-control O.D. value) ×100. Sera with percent inhibition values of at least 20% were regarded as positive [14]. For samples collected until November 10, 2020, sera were tested by both VN and sVNT in parallel. Subsequently, sera were screened by sNVT first, and only positive samples were tested by VN.

### Criteria for determining animal infection by SARS-CoV-2

Any animal that had oral, nasal or respiratory swab samples where viral RNA was detected by RT-qPCR for at least two genes of SARS-CoV-2 with Ct values ≤40 was considered actively infected with SARS-CoV-2 at the time of sampling. Pets that had only body (fur) swabs with Ct values ≤40 were considered negative for SARS-CoV-2 infection. Animals that had any serum sample presenting neutralizing antibodies via VN were considered previously exposed to SARS-CoV-2. Both currently infected and/or previously exposed animals constitute cases of animal infection with SARS-CoV-2.

### Risk factors for SARS-CoV-2 infection

To assess putative risk factors associated with infection, project data were categorized into the following: sampling aspects, household characteristics, animal aspects and human aspects. Sampling aspects included number of days between the human diagnosis of COVID-19 and animal sampling. Household characteristics included number of pets per household and number of people with known COVID-19 in the household at the time of animal sampling. For that, we compared households with one versus two or more people with COVID-19. Animal aspects included age, sex, species, sterilization, travel history, and access to outdoors, assigning as pets without access to outdoors when reported to spend 90% or more of their time indoors. Human aspects included the pet’s level of contact with people, such as whether people with COVID-19 shared a room, bed, and food with the pet, as well as petting and kissing/licking during the self-isolation.

### Statistical analysis

We used either Chi-squared test with Yates’ correction or Fisher’s Exact test to compare positivity by RT-qPCR, VN and SARS-CoV-2 infection between dogs and cats. We compared endpoint titers from VN testing between dogs and cats. These values were not normally distributed (assessed using Shapiro-Wilk normality test, data not shown), and, therefore, were compared using Wilcoxon signed-rank test. This same test was employed to evaluate whether there were differences in the number of days between human diagnosis of COVID-19 and sampling for dogs and cats.

We built logistic regression models to determine factors associated with the risk of dogs and cats being positive by either RT-qPCR or seropositive by VN separately. For these models, we included results from the first sampling event only. The initial model included number of days after diagnosis, pet sex, age and sterilization status, number of pets in the household, pet access to outdoors, recent travel, and whether the household had one or more than one person with COVID-19. For factors related to whether infected people interacted with the pets during quarantine, we detected high multicollinearity (correlation values above or below 0.85 and −0.85, respectively). Therefore, we only included sleeping in the same room, sharing food, and petting as explanatory variables for our models. We performed backward stepwise selection using Akaike Information Criterion (AIC) to identify significant predictors in these models. Coefficients from the final models were used to calculate odds ratios (OR) and their 95% confidence intervals (CI). We performed all analyses using R 4.2.2 [15].

## Results

### General results

Between June 13, 2020, and May 18, 2021, we sampled 579 companion animals from 281 households in ten counties in Texas. This included 396 (71%) dogs, 157 (29%) cats, and 26 (4.5%) individuals of ten other species, namely Guinea pigs (n=6), horses (n = 5), rabbits (n = 4), ferrets (n = 2), rats (n = 2), alpaca (n = 2), a hamster, a hedgehog, bearded dragons (n = 2), and a parrot. The median number of pets per household was 2, ranging from one to 13 pets (Figure 1). The mean time from human diagnosis to animal sampling was 12.1 days (SD = 12.0; 95% CI: 10.9–13.3) for dogs and 11.1 days (SD = 10.1; 95% CI: 9.56–12.7) for cats, with no significant difference between species (*P* = 0.67).

**Figure 1.**
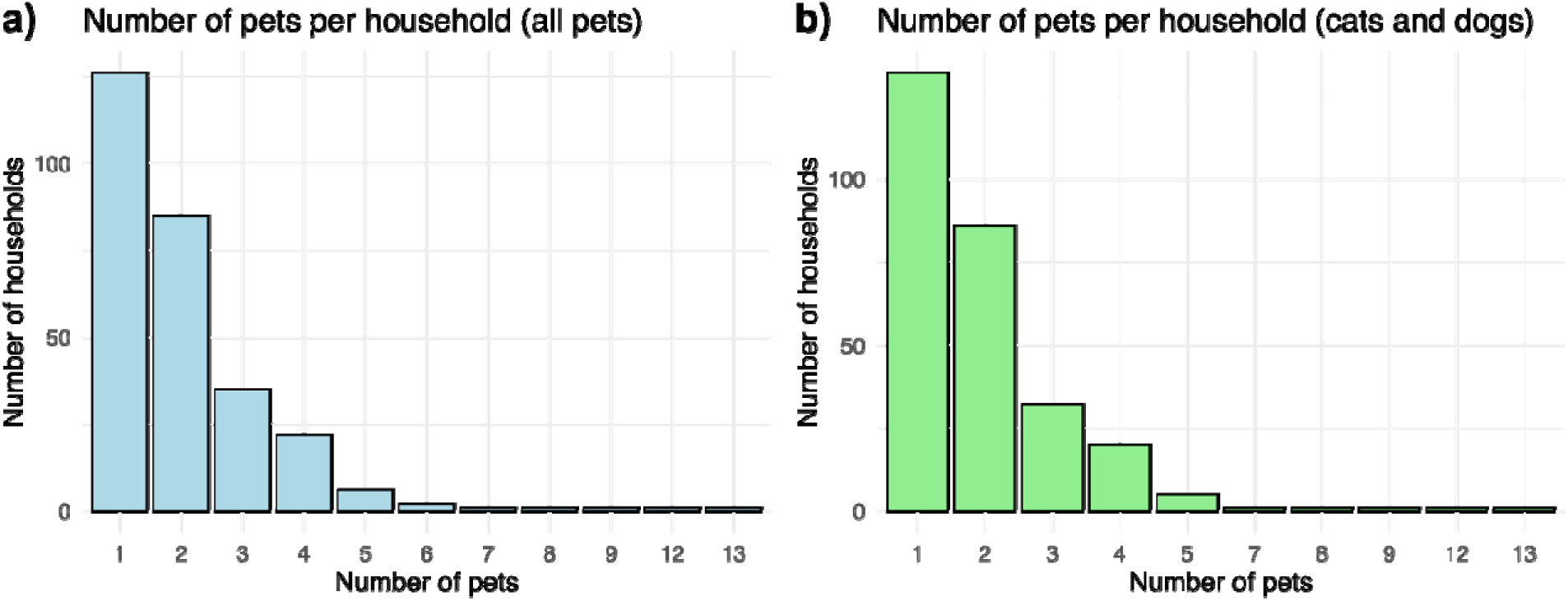
Distribution of the number of pets per household sampled between June 13, 2020, and May 18, 2021, in central Texas, USA. a) distribution across all sampled pets, including dogs, cats, and other species, and b) distribution of only cats and dogs.

SARS-CoV-2 RNA was detected in 4.8% (19/396) of the dogs and 13.4% (21/157) of the cats tested by RT-qPCR. A total of 56 swab samples from 40 animals tested positive for SARS-CoV-2 by RT-qPCR (Supplemental material Table S1). Nine (20%) animals tested positive for SARS-CoV-2 by RT-qPCR for body swabs only and were not considered as infected. When comparing swab types that were collected simultaneously, nasal swabs were most frequently positive (n = 29 animals). Notably, body swab had the second highest number of positives (n = 25 animals) followed by oral (n = 20), rectal (n = 7) and conjunctival (n=2).

Neutralizing antibodies were detected in 24.9% (95/382) of the dogs and 35.6% (52/146) of the cats tested. The endpoint titers from the VN testing were significantly higher for cats than dogs (Wilcoxon test, *P* < 0.001; Figure 2). In cats, the titer range was 8 to 2,048 with a mean VN titer of 182.14 when considering results from the first sampling event. In dogs, the titer range was 8 to 128, with a mean VN titer of 34.8.

**Figure 2.**
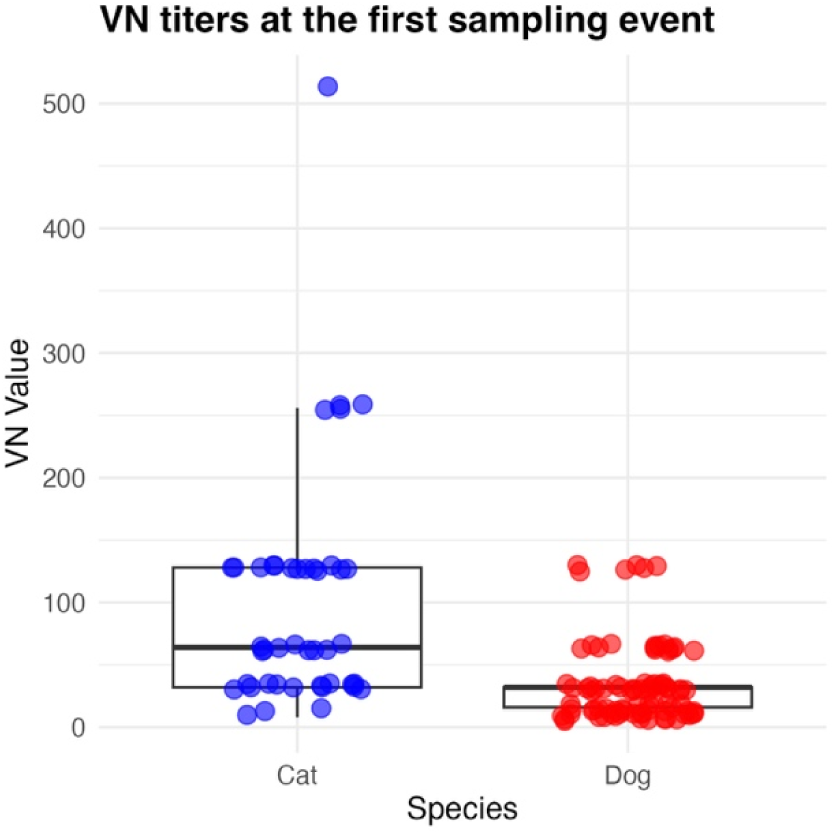
Virus neutralization (VN) titers for dogs and cats at the first sampling event. VN titers are represented as boxplots with individual data points and are shown as the reciprocal of th highest serum dilution that provided full neutralization of SARS-CoV-2. Median, interquartile range (IQR), and whiskers representing 1.5 times the IQR are displayed.

Detection frequency was lower in dogs compared to cats by RT-qPCR (χ2 with Yates correction = 11.1; *P* < 0.001) and by VN (χ2 with Yates correction = 5.5; *P* = 0.018). However, infection rates, based on combined RT-qPCR and VN results, did not differ between dogs (108/396; 27.3%) and cats (56/157; 35.7%; χ2 with Yates correction = 3.4, *P* = 0.064). None of the other 10 species tested positive for SARS-CoV-2 RNA or for neutralizing antibodies. In total, 110 out of 281 (39.1%) households across six counties had at least one pet positive for SARS-CoV-2 infection by RT-qPCR or serology.

During the first sampling event, pets were sampled between 0 and 86 days after the initial COVID-19 diagnosis in household members, with a median of 8 days. The median number of days after diagnosis for RT-qPCR-positive dogs was 7 days (SD = 3.37; 95% CI: 4.74–7.99), while for cats it was 6 days (SD = 4.33; 95% CI: 5.32–9.25). Among seropositive pets, the median number of days after diagnosis for dogs was 11 days (SD = 14.1; 95% CI: 12.6–18.6), and for cats, it was 13 days (SD = 13.4; 95% CI: 11.4–19.6). These values did not change between species (*P* > 0.05) No pet tested positive for RT-qPCR when sampled on or more than 18 days after COVID-19 diagnosis, although seropositivity was detected when the initial sample was taken up to 86 days after the human diagnosis.

Longitudinal sampling occurred with 28 dogs, 30 cats, and two horses from 28 households, ranging from two to eight sampling events per animal. Among pets sampled more than once, eight dogs and 14 cats were RT-qPCR-positive, and 17 dogs and 27 cats had neutralizing antibodies against SARS-CoV-2 in at least one sampling event (Supplemental material Figure S1). In every case, RT-qPCR-positivity was detected during the first sampling event, with two cats also testing positive in the second event. Seropositivity was identified in eight dogs and 18 cats during the first sampling event and seroconversion occurred during the second sampling event for nine dogs and nine cats. Throughout our study, three cats and 10 dogs remained seronegative despite being resampled and living in the same house with at least one other positive animal.

### Factors associated with SARS-CoV-2 infection in dogs and cats

When considering results from the first sampling event only, we identified factors associated with increased odds of infection by SARS-CoV-2 in dogs (Table 1) and cats (Table 2). Chances of detecting viral RNA decreased in dogs (OR = 0.90, CI: 0.8–0.97, *P* = 0.032) and in cats (OR = 0.90, CI: 0.8–0.98, *P* = 0.023) as number of days after human diagnosis progressed; each extra day is associated with a 10% decrease in the chances of dogs and cats testing positive. Conversely, each additional day increased the odds of seropositivity by 2% for dogs and 5% for cats. Additionally, older dogs had a 13% increase in odds of testing positive by RT-qPCR for every additional year of age (OR = 1.13, 95% CI: 1–1.26, *P* = 0.036). Seropositivity was associated with sharing food during self-isolation, increasing more than five-fold for cats (OR = 5.50, 95% CI: 1.72–19.57, *P* = 0.005) and two-fold for dogs (OR = 2.06, 95% CI: 1.12–3.73, *P* = 0.018). Seropositivity also increased with time from human diagnosis in dogs (OR = 1.02, 95% CI: 1–1.04, *P* = 0.027) and cats (OR = 1.05, 95% CI: 1–1.1, *P* = 0.035). Sleeping in the same room as household members during self-isolation increased chances for dogs becoming seropositive (OR = 2.34, 95% CI: 1.36–4.13, *P* = 0.005). Additionally, living in households with two or more people with COVID-19 was correlated with more than a two-fold increase in seropositivity in dogs compared to those living in households with a single person with COVID-19 (OR = 2.51, 95% CI: 1.47–4.39, *P* = 0.001).

**Table 1.**
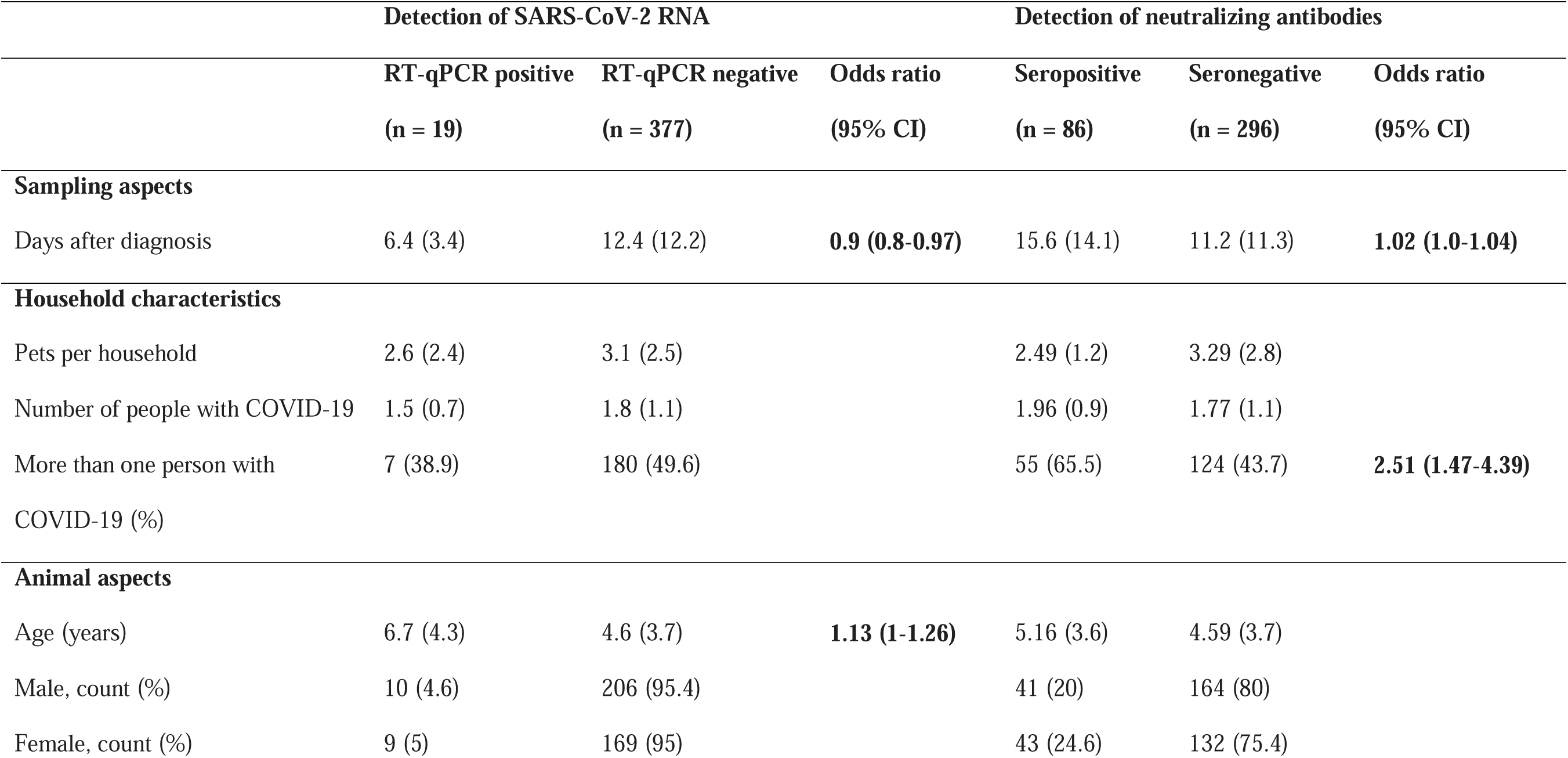
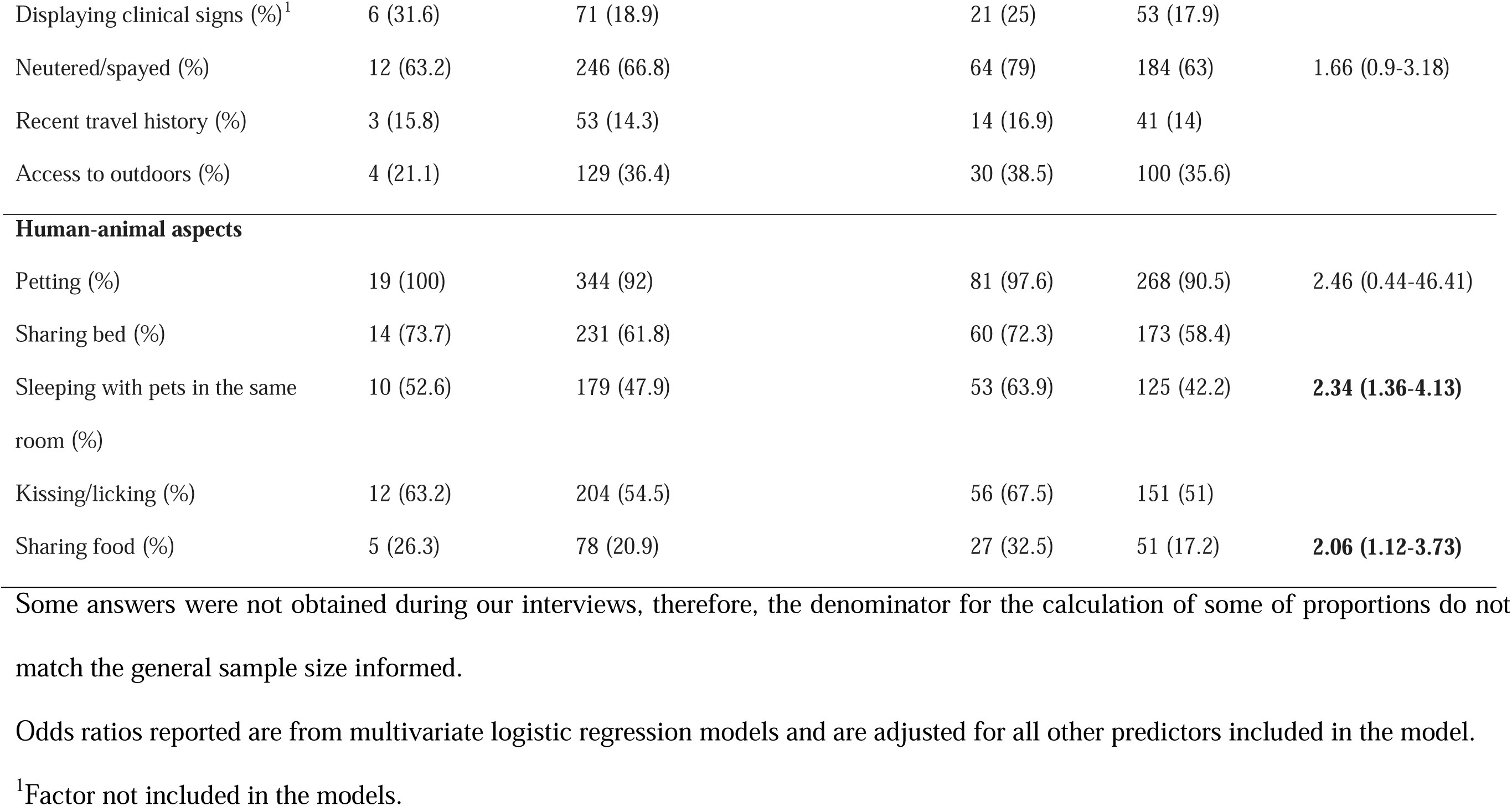
Risk factors for SARS-CoV-2 infection in dogs from households with people reporting COVID-19 in Texas. Logistic regression models were used to determine factors associated with detection of viral RNA and neutralizing antibodies against SARS-CoV-2. Values are mean followed by standard deviation (in parenthesis) unless otherwise stated. Adjusted odds ratio and 95% confidence intervals (CI) are reported for factors kept in the final models; statistically significant values are in bold.

**Table 2.**
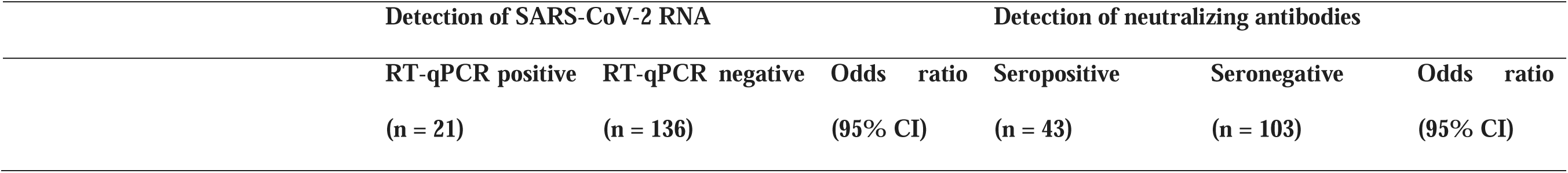
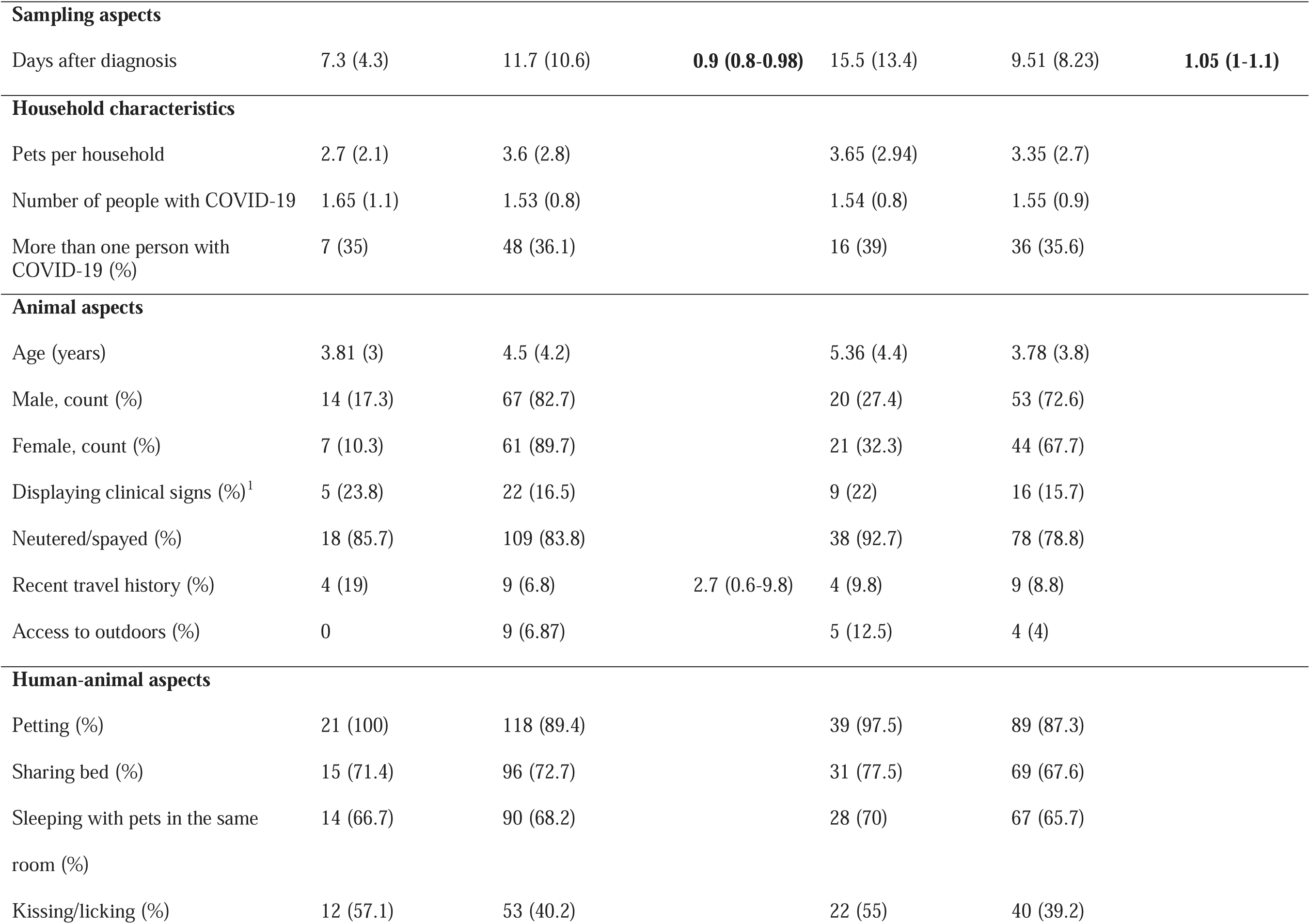
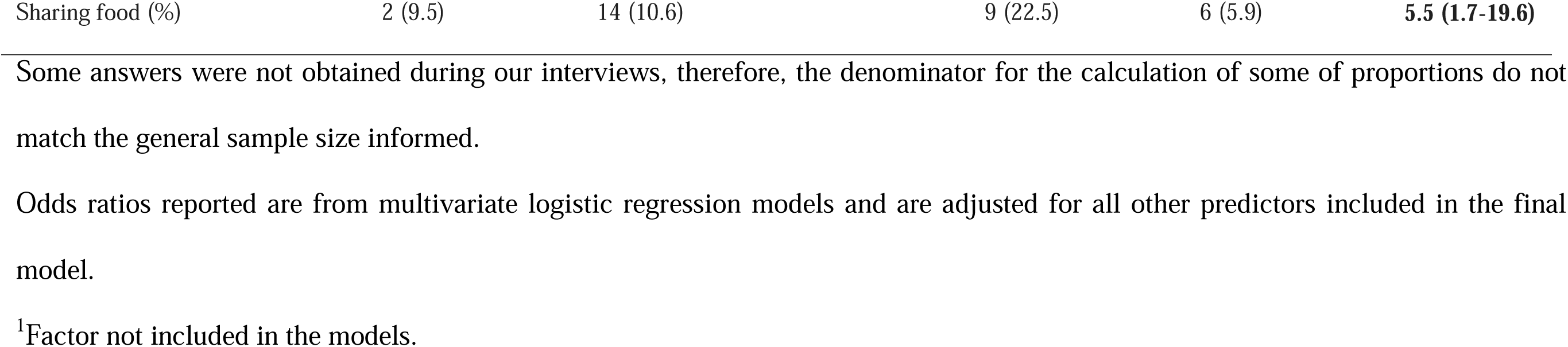

### Clinical signs of disease

Frequency of reported clinical signs did not differ between infected and non-infected pets (χ2 with Yates correction; *P* > 0.05; Figure 3). Among dogs positive by RT-qPCR, 31.6% (6/19) displayed clinical signs, while 18.9% (71/375) of RT-qPCR-negative dogs displayed clinical signs. Similarly, frequency of clinical signs for RT-qPCR positive and negative cats were 23.8% (5/21) and 16.5% (22/133), respectively. According to serological data, 28.4% (21/74) seropositive dogs had clinical signs, while 20.6% (63/306) seronegative dogs displayed clinical signs. Frequencies of clinical signs observed in seropositive and seronegative cats were 36% (9/25) and 27.1% (32/118), respectively. The proportion of pets displaying clinical signs detailed by category (respiratory, gastrointestinal or behavioral) according to positivity for SARS-CoV-2 is presented in Figure 3.

**Figure 3.**
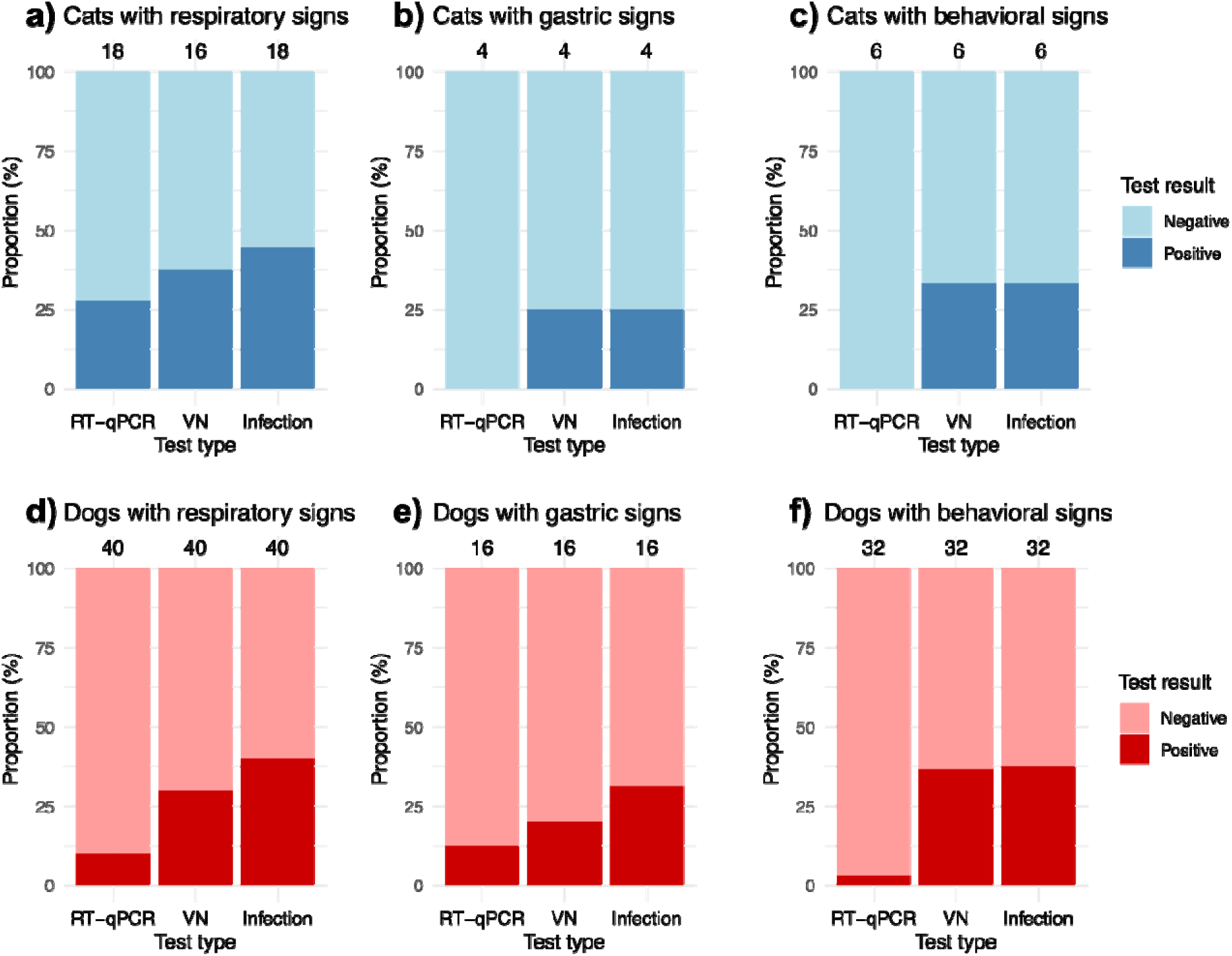
Owner-reported clinical signs in pets living in COVID-19 positive households, Texas, 2020-2021. Pets are stratified into those that tested negative and positive for SARS-CoV-2 in our study. Panels a, b, and c represent cats with respiratory, gastric, and behavioral signs, respectively. Panels d, e, and f correspond to dogs with the same categories of clinical signs. Results are displayed for RT-qPCR, VN, and infection status (RT-qPCR or VN positive). Sample sizes are shown above the bars. The prevalence of clinical signs was not statistically different between SARS-CoV-2 infected versus uninfected animals.

### Virus isolation and whole genome sequencing

We isolated SARS-CoV-2 from respiratory, oral or nasal samples collected from six pets (Table 3) from a total of 19 that had samples RT-qPCR-positive samples submitted for virus isolation (Supplemental material Table S2). Whole genome sequencing was successful for 8 dogs and 16 cats from respiratory, oral and nasal swabs, and in the case of one cat, a rectal swab, revealing several lineages, including the VOC Alpha (B.1.1.7), and B.1.234, B.1.1, B.1, and B.1.609 (Table 4).

**Table 3.** SARS-CoV-2 isolates confirmed by cytopathic effect in Vero cell cultures, followed by RT-qPCR and nucleotide sequencing from respiratory samples collected from cats and dogs from Texas, United States, 2020-2021. Samples 466 and 467 were reported previously [8].

| Animal ID | Species | Swab | Date of collection | RT-qPCR (Ct values)<br>RdRp/E/N1-N2 | GISAID | Pango Lineage | Clade | Isolate ID | Vero 76 culture (passage) |
| --- | --- | --- | --- | --- | --- | --- | --- | --- | --- |
| TAMU-013 | Cat | R | 6/28/2020 | 21.73/21.12/18-17.7 | EPI_ISL_699506 | B.1.234 | G | hCoV-19/cat/USA/TX-TAMU-013/2020 | (2) |
| TAMU-466 | Dog | O | 2/12/2021 | NT/NT/28.2-28.5 | EPI_ISL_1315074 | B.1.1.7 | GRY | hCoV-19/dog/USA/TX-TAMU-21-005988-002-466/2020 | (2) |
| TAMU-172 | Cat | N | 9/11/2020 | NT/NT/27.8-27.4 | EPI_ISL_6639377 | B.1.1 | GR | hCoV-19/cat/USA/TX-TAMU-172-20-027713-002/2020 | (3) |
| TAMU-252 | Cat | O | 10/15/2020 | NT/NT/27.5-27.8 | EPI_ISL_3010055 | B.1 | GH | hCoV-19/cat/USA/TX-TAMU-252-20-031373-005/2020 | (3) |
| TAMU-392 | Dog | N | 12/22/2020 | NT/NT/29.6-29.8 | EPI_ISL_3148877 | B.1.609 | G | hCoV-19/dog/USA/TX-TAMU-392-21-002748-002/2020 | (1) |
| TAMU-467 | Cat | N | 2/12/2021 | NT/NT/24.6-24.6 | EPI_ISL_1315075 | B.1.1.7 | GRY | hCoV-19/cat/USA/TX-TAMU-21-005988-005-467/2021 | (2) |

R - Respiratory, includes nasal and oral swabs together; O - Oral; N - Nasal; RdRp - RNA-dependent RNA polymerase gene; E - Envelope small membrane protein gene; N1 and N2 - regions of the nucleocapsid gene; GISAID - Global Initiative on Sharing All Influenza Data; Ct - Cycle threshold,

**Table 4.**
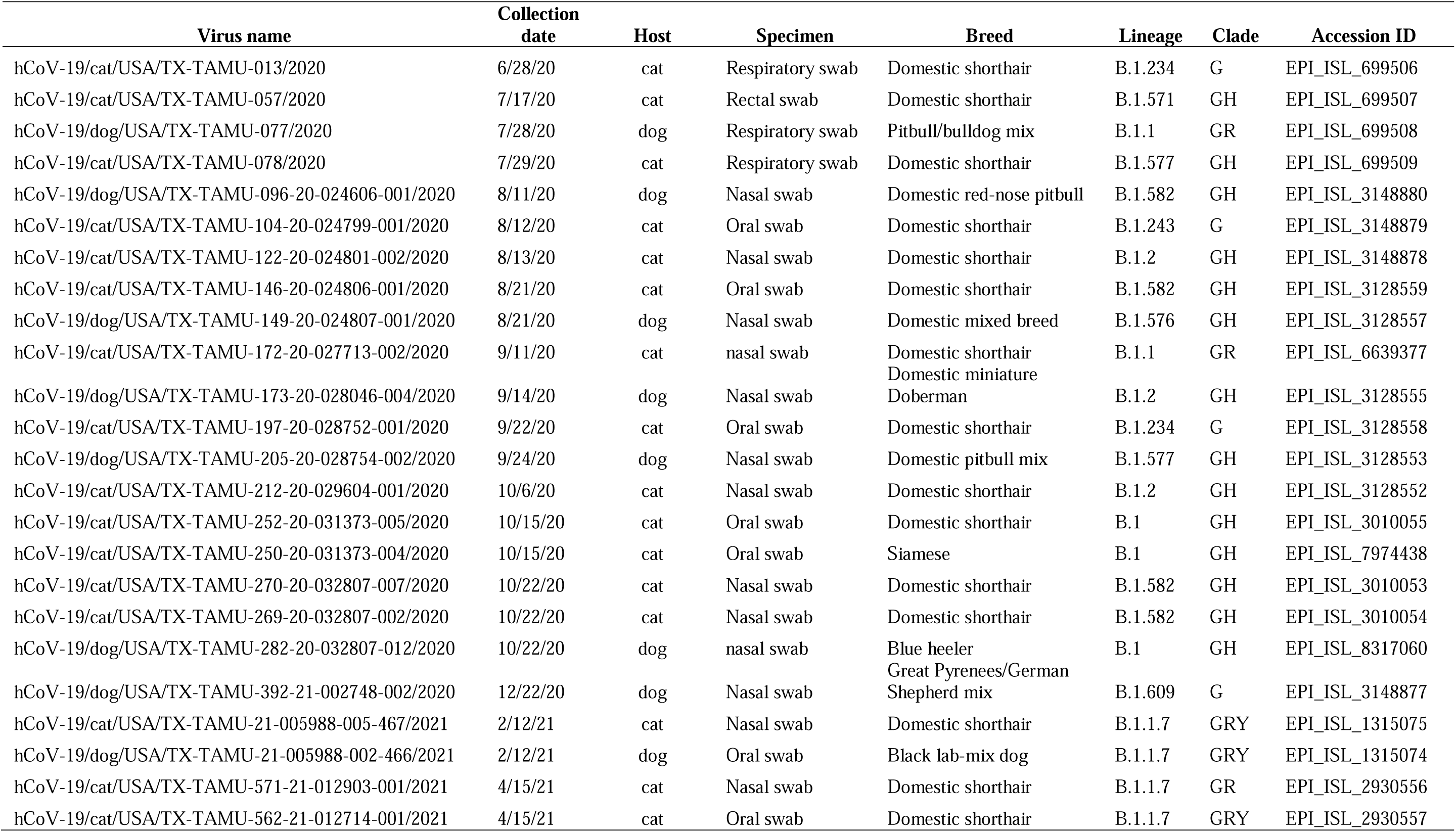
SARS-CoV-2 viral genome sequences from cats and dogs from Texas, United States, 2020-2021.

## Discussion

Our study showed that active surveillance of pets in high-risk environments can provide key information on the frequency and timing of animal infections in relation to human infections. Roughly one in every four companion animals (164 of 579; 28.8%) that lived with at least one person with COVID-19 was confirmed to have an active or past SARS-CoV-2 infection, based on active household surveillance from June 2020 to May 2021 in East-Central Texas, USA. With respect to households, 110 of 281 (39%) contained pets that had been infected with SARS-CoV-2. The surveillance conducted in this study is the largest reported in terms of households with confirmed COVID-19 in people being investigated for zoonotic transmission through spillback to companion animals. This work led Texas being placed as the state with the highest estimated number of confirmed cases of SARS-CoV-2 infection in pets in the USA [16]. In our study, only dogs and cats were positive for SARS-CoV-2 among 12 species of companion animals investigated, but the sample sizes for other species were low. Lizards and birds were sampled to include all pets from households that also had cats and/or dogs, but are not known to be part of the natural range of SARS-CoV-2 hosts. Infections in dogs and cats have been reported worldwide [7,17–19], but the documented exposure of other species of companion animals remains limited. The higher prevalence for SARS-CoV-2 RNA and neutralizing antibodies in cats during the first stages of the COVID-19 pandemic found here corroborates previous studies in multiple countries [17,20,21].

The odds of detecting SARS-CoV-2 RNA in dogs and cats decreased with increasing days post-human COVID-19 diagnosis. Current evidence suggests SARS-CoV-2 RNA is detected for up to 15 days in asymptomatic persons and for 30 days after symptom onset in mildly and severely affected groups [22]. Rapid active surveillance sampling animals with a median of 11-12 days following human diagnosis likely resulted in the high apparent frequency of active animal infections, for which the median days of SARS-CoV-2 detection was 7 days for dogs and 6 days for cats. In a passive surveillance study, the median number of days from presumed exposure to detection of active infections was of 6 days in dogs of 10 days in cats [23], further supporting the importance of early sampling to maximize detection likelihood.

We found that prevalence of neutralizing antibodies increased with the number of days since human diagnosis. Serum neutralization assays represent the gold standard for assessing antibody-mediated protection in naturally infected and vaccinated individuals for SARS-CoV-2 [24]. The use of a gold standard method is also crucial for a reliable investigation of the diverse group of coronaviruses reducing the chance of misclassification of an animal as seropositive.

Few pet-related and human-pet interaction factors explained risks of infections in dogs and cats. Sharing food was a significant risk factor for seropositivity in both dogs and cats, while sleeping in the room with pets was a significant risk factor for dogs only. In contrast, in a household transmission study in Canada, Bienzle et al, [5] observed that cats, but not dogs, had a higher likelihood of SARS-CoV-2 seropositivity when they slept in their owner’s bed. The difference observed between the two species may be related to their different behavior, and perhaps difference in susceptibility related to ACE2. We observed that cats were just as likely to be infected regardless of whether there was very close contact with humans (e.g., sharing a bed) or not, possibly because of differences in biological susceptibility [25]. Given several significant risk factors identified through this study that relate to close interactions between people and pets during outbreaks or pandemics, it is critical for public health and animal health officials to provide recommendations to prevent transmission of zoonotic diseases. Overall, our results support the notion that people with COVID-19 should also isolate from animals to reduce the risk of SARS-CoV-2 infection in pets.

Seroprevalence among dogs living in households with two or more people with COVID-19 was significantly higher than among dogs from households with a single person with COVID-19. However, this factor did not influence SARS-CoV-2 RNA detection in dogs, nor was it associated with higher odds of infection for cats. These findings may reflect higher susceptibility of cats, which could result in similar exposure regardless of the number of infected people.

We found that signs of illness were not significantly more frequent among infected than uninfected animals. Therefore, reported clinical signs could relate to background illness (not related to SARS-CoV-2) or to a closer level of observation by owners of their pets during their time of self-isolation. These findings are similar to a serological study assessing 1000 pets across the United States [26]. Alternately, some apparently uninfected animals with reported clinical signs may have been infected. The clinical signs recorded here were described by the owners through the questionnaire, and no clinical evaluation was done by a veterinarian during sample collections. Experimental infection studies have demonstrated that despite being susceptible to infection by SARS-CoV-2, cats do not develop clear clinical signs [27,28]. Another experimental infection study demonstrated that the respiratory tract of cats is more severely affected by SARS-CoV-2 than that of dogs [29]. Field investigations have found evidence of illness, related or not, in naturally SARS-CoV-2-infected companion animals worldwide [7,17,20,30].

Despite evidence of companion animals being infected by SARS-CoV-2 worldwide, transmission from companion animals to humans remains rare [31]. A single case of SARS-CoV-2 transmission from a cat to a human in Thailand was reported in August 2021 when a veterinarian was sneezed on by an exposed, asymptomatic cat [32]. In our investigation, the isolation of virus in cell culture from multiple pets suggests that pets may be infectious, but onward transmission from animals to people was not measured in this study. We isolated multiple lineages of SARS-CoV-2 from respiratory swabs of six animals (both cats and dogs), including Alpha, B.1.234, B.1.1, B.1, and B.1.609, indicating potential capacity to act as source of infection for their contacts. Virus isolation of SARS-CoV-2 from companion animals has not been reported frequently, as most diagnoses are made by RT-qPCR or serological kits later confirmed in reference laboratories [33].

The SARS-CoV-2 lineages previously reported in the human populations and isolated from companion animals described here likely represent zoonotic transmission through spillback from the owners or community to the animals. Lineage B.1.234 was isolated from cat TAMU-013 on June 28^th^, 2020, as the only RT-qPCR-positive pet in a household that also had two dogs. Cat TAMU-13 was sampled only 11 days after the first collection date of this lineage in humans in Texas [34]. Lineage B.1.1 was isolated from the nasal swab of cat TAMU-172 three days after human diagnosis. In Texas, the cumulative prevalence of this lineage was 1% among all SARS-CoV-2 sequenced uploaded into GISAID [35]. TAMU-172 was seven years-old, lived in a household with no other companion animal, and had history of sharing bed with the owner during self-isolation. Lineage B.1 was isolated from a cat sampled six days after human diagnosis in mid-October 2020. The cat TAMU-252 was four months-old and had history of sharing bed with the owner. This lineage reached an average daily prevalence of over 60% in the United States between March and April 2020 [34]. Lineage B.1.609 was isolated from a four-year old dog (TAMU-392) that was sampled a week after human diagnosis in a household with two people with COVID-19 and five other companion animals. TAMU-392 shared the bed with the owner and was the only positive companion animal in the household. This lineage was first detected in Texas in May 2020 and had a cumulative prevalence of less than 0.5%. Finally, we previously detailed the first detection of the Alpha variant B.1.1.7 in companion animals in the US; this occurred in a pet dog and cat that lived together [8].

### Conclusion

Pet infections were not uncommon in households where an infected human resided, with roughly 30% of dogs and cats having SARS-CoV-2 infection in 39% of the 281 households sampled. Cats were more likely to be positive by RT-qPCR and by virus neutralization test than dogs. Given the median number of days elapsed from the day of human COVID-19 diagnosis to sample collection from household pets that ultimately tested RT-qPCR-positive for SARS-CoV-2 was only 6-7 days, we suggest that future pet surveillance studies in such high-risk environments prioritize sample collection within a week of human diagnosis. Sharing food and sleeping in the same room with infected people were associated with increased risk for companion animal infection in households with people with COVID-19. Therefore, preventive measures including the restriction or at least the reduction of close contact between people with COVID-19 and pets protect pets from SARS-CoV-2 infection. Despite reports of symptomatic pets testing positive for SARS-CoV-2, our study shows that infected pets were not more likely to have clinical signs than uninfected pets, and so passive surveillance pipelines that rely on testing of symptomatic animals only may be less useful than active surveillance. Additionally, we showed that pets can signal viral lineages contemporaneously present in human populations, sometimes at nearly the same time as the first human reports of distinct lineages, emphasizing that genomic surveillance efforts should also include pets. Characterizing animal infections using active SARS-CoV-2 surveillance in pets at risk of infection using the One Health approach is a critical step to effectively address pandemic prevention, response, and management.

## Acknowledgements

We thank our partners at the Centers for Disease Control and Prevention for collaboration, and all pet owners for their cooperation. This research was funded by Texas A&M AgriLife Research and the Centers for Disease Control and Prevention RFP 75D 301-20-R-68167. The findings and conclusions in this report are those of the authors and do not necessarily represent the official position of the Centers for Disease Control and Prevention.

---

**Supplemental material Figure S1.**
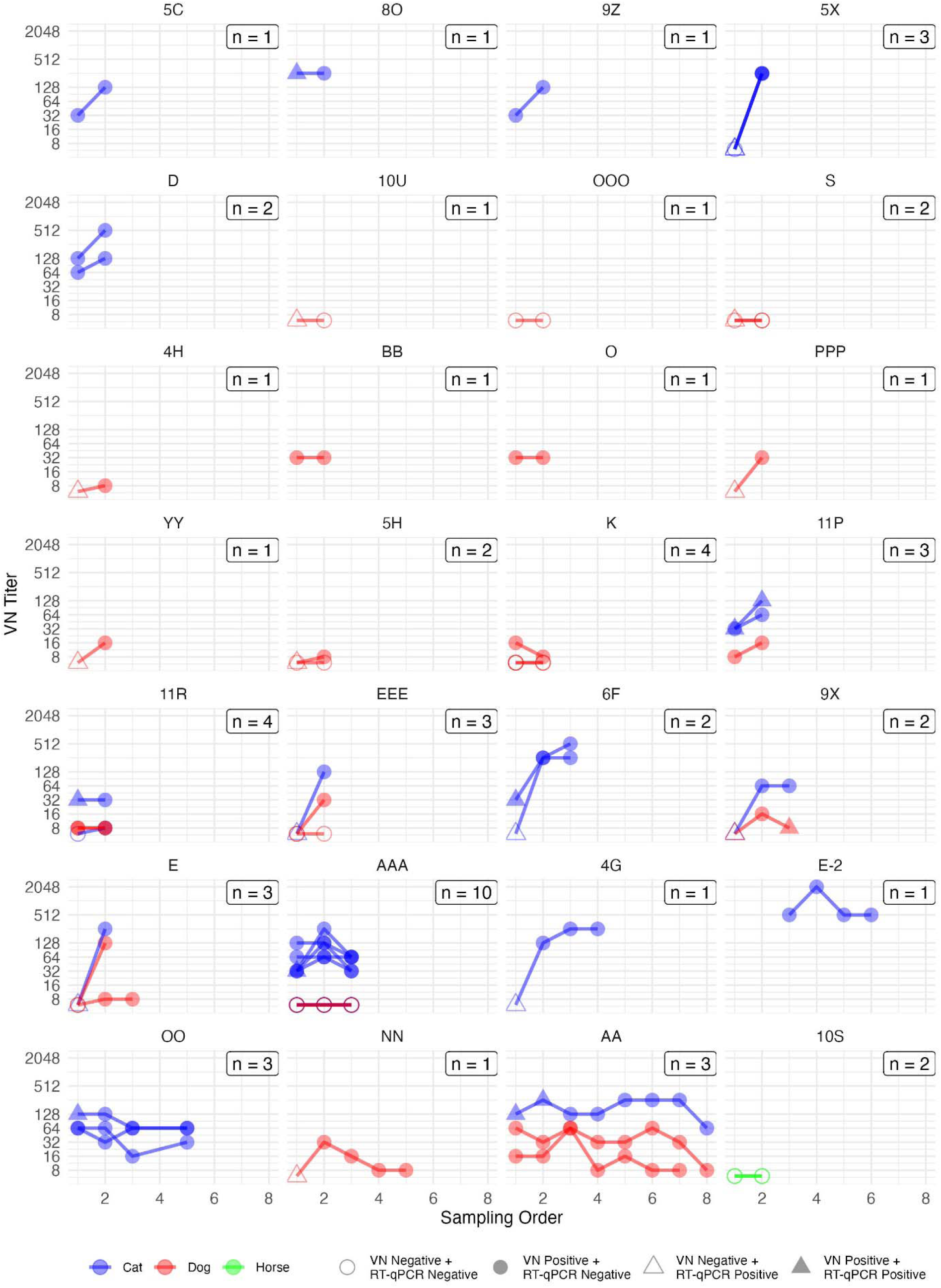
Longitudinal sampling among pets in COVID-19 positive households, Texas, 2020-2021. The pets in households with at least one animal found to be positive at the initial sampling event in all but two households were re-sampled over time, with up to eight total sampling events. Each household is plotted separately, with 1-10 pets per house (number of pets are shown inside boxes for each household). RT-qPCR positivity was restricted to the first one to two sampling events only, whereas seropositivity was detected in some animals throughout the study period with generally stable or increasing antibody titers, including seroconversion in some animals.

Supplemental material Table S1. Cycle threshold values of swabs of dogs and cats that tested positive for SARS-CoV-2 by RT-qPCR in Texas, United States. ID-Animal identification; HH-Household identification; HD-Human diagnostic; CD-Sample collection date; DAHD-Days after human diagnosis; TAMU-Texas A&M University Laboratory; WVDL-Wisconsin Veterinary Diagnostic Laboratory; NVSL-National Veterinary Services Laboratories; RdRp-RNA-dependent RNA polymerase gene; E-Envelope gene; N1-Nucleocapsid gene region 1; N2-Nucleocapsid gene region 2; ND-Not detected; Squares with no Ct values indicate not tested; Samples B and C indicate sequential samples. Gray shaded rows show animals that were re-sampled with RT-qPCR positive results.

Supplemental material Table S2. RT-qPCR-positive swab samples submitted to virus isolation but that did not produce cytopathic effects in Vero cell cultures.

## Uncategorized References

1. [1] Davies NG, Jarvis CI, Group CC-W, et al. Increased mortality in community-tested cases of SARS-CoV-2 lineage B.1.1.7. Nature. 2021 May;593(7858):270–274.

2. [2] Sanyaolu A, Okorie C, Marinkovic A, et al. The emerging SARS-CoV-2 variants of concern. Therapeutic Advances in Infectious Disease. 2021 2021-01-01;8:204993612110243.

3. [3] World Organisation for Animal Health - SARS-COV-2 in animals - situation report 22 [Internet]. WOAH. 2023 [cited 30/06/2023].

4. [4] Ghai RR, Carpenter A, Liew AY, et al. Animal Reservoirs and Hosts for Emerging Alphacoronaviruses and Betacoronaviruses. Emerg Infect Dis. 2021 Apr;27(4):1015–1022.

5. [5] Bienzle D, Rousseau J, Marom D, et al. Risk factors for SARS-CoV-2 infection and illness in cats and dogs. Emerging Infectious Diseases. 2022;28(6):1154.

6. [6] Kannekens-Jager MM, de Rooij MMT, de Groot Y, et al. SARS-CoV-2 infection in dogs and cats is associated with contact to COVID-19-positive household members. Transbound Emerg Dis. 2022 Nov;69(6):4034–4040.

7. [7] Hamer SA, Pauvolid-Correa A, Zecca IB, et al. SARS-CoV-2 Infections and Viral Isolations among Serially Tested Cats and Dogs in Households with Infected Owners in Texas, USA. Viruses. 2021 May 19;13(5).

8. [8] Hamer SA, Ghai RR, Zecca IB, et al. SARS-CoV-2 B.1.1.7 variant of concern detected in a pet dog and cat after exposure to a person with COVID-19, USA. Transbound Emerg Dis. 2022 May;69(3):1656–1658.

9. [9] Corman VM, Landt O, Kaiser M, et al. Detection of 2019 novel coronavirus (2019-nCoV) by real-time RT-PCR. Euro Surveill. 2020 Jan;25(3).

10. [10] Konrad R, Eberle U, Dangel A, et al. Rapid establishment of laboratory diagnostics for the novel coronavirus SARS-CoV-2 in Bavaria, Germany, February 2020. Euro Surveill. 2020 Mar;25(9).

11. [11] Lu X, Wang L, Sakthivel SK, et al. US CDC Real-Time Reverse Transcription PCR Panel for Detection of Severe Acute Respiratory Syndrome Coronavirus 2. Emerg Infect Dis. 2020 Aug;26(8).

12. [12] Goryoka GW, Cossaboom CM, Gharpure R, et al. One Health investigation of SARS-CoV-2 infection and seropositivity among pets in households with confirmed human COVID-19 cases—Utah and Wisconsin, 2020. Viruses. 2021;13(9):1813.

13. [13] McAloose D, Laverack M, Wang L, et al. From People to Panthera: Natural SARS-CoV-2 Infection in Tigers and Lions at the Bronx Zoo. mBio. 2020 Oct 13;11(5).

14. [14] Perera R, Ko R, Tsang OTY, et al. Evaluation of a SARS-CoV-2 Surrogate Virus Neutralization Test for Detection of Antibody in Human, Canine, Cat, and Hamster Sera. J Clin Microbiol. 2021 Jan 21;59(2).

15. [15] Team RC. R: A language and environment for statistical computing. R Foundation for Statistical Computing, Vienna, Austria. https://www.R-project.org/. 2022.

16. [16] SARS-CoV-2 in Animals [Internet]. Animal and Plant Health Inspection Service, U.S. Department of Agriculture: 2024 [cited 19/12/2024].

17. [17] Calvet GA, Pereira SA, Ogrzewalska M, et al. Investigation of SARS-CoV-2 infection in dogs and cats of humans diagnosed with COVID-19 in Rio de Janeiro, Brazil. PLoS One. 2021;16(4):e0250853.

18. [18] Klaus J, Zini E, Hartmann K, et al. SARS-CoV-2 Infection in Dogs and Cats from Southern Germany and Northern Italy during the First Wave of the COVID-19 Pandemic. Viruses-Basel. 2021 Aug;13(8).

19. [19] Imanishi I, Asahina R, Hayashi S, et al. Serological Study of SARS-CoV-2 Antibodies in Japanese Cats: Analysis of Risk Factors Among Cat Lifestyles. Research Square Platform LLC; 2022.

20. [20] Sailleau C, Dumarest M, Vanhomwegen J, et al. First detection and genome sequencing of SARS-CoV-2 in an infected cat in France. Transbound Emerg Dis. 2020 Nov;67(6):2324–2328.

21. [21] Ruiz-Arrondo I, Portillo A, Palomar AM, et al. Detection of SARS-CoV-2 in pets living with COVID-19 owners diagnosed during the COVID-19 lockdown in Spain: A case of an asymptomatic cat with SARS-CoV-2 in Europe. Transbound Emerg Dis. 2021 Mar;68(2):973–976.

22. [22] Kim DY, Bae EK, Seo JW, et al. Viral Kinetics of Severe Acute Respiratory Syndrome Coronavirus 2 in Patients with Coronavirus Disease 2019. Microbiol Spectr. 2021 Oct 31;9(2):e0079321.

23. [23] Liew AY, Carpenter A, Moore TA, et al. Clinical and epidemiologic features of SARS-CoV-2 in dogs and cats compiled through national surveillance in the United States. Journal of the American Veterinary Medical Association. 2023 02 Jan. 2023:1–10.

24. [24] Matusali G, Colavita F, Lapa D, et al. SARS-CoV-2 Serum Neutralization Assay: A Traditional Tool for a Brand-New Virus. Viruses. 2021 Apr 10;13(4).

25. [25] Stout AE, Andre NM, Jaimes JA, et al. Coronaviruses in cats and other companion animals: Where does SARS-CoV-2/COVID-19 fit? Vet Microbiol. 2020 Aug;247:108777.

26. [26] Kimmerlein AK, McKee TS, Bergman PJ, et al. The Transmission of SARS-CoV-2 from COVID-19-Diagnosed People to Their Pet Dogs and Cats in a Multi-Year Surveillance Project. Viruses. 2024;16(7):1157.

27. [27] Bosco-Lauth AM, Hartwig AE, Porter SM, et al. Experimental infection of domestic dogs and cats with SARS-CoV-2: Pathogenesis, transmission, and response to reexposure in cats. P Natl Acad Sci USA. 2020 Oct 20;117(42):26382–26388.

28. [28] Halfmann PJ, Hatta M, Chiba S, et al. Transmission of SARS-CoV-2 in Domestic Cats. N Engl J Med. 2020 Aug 6;383(6):592–594.

29. [29] Shi J, Wen Z, Zhong G, et al. Susceptibility of ferrets, cats, dogs, and other domesticated animals to SARS-coronavirus 2. Science. 2020 May 29;368(6494):1016–1020.

30. [30] Hosie MJ, Epifano I, Herder V, et al. Detection of SARS-CoV-2 in respiratory samples from cats in the UK associated with human-to-cat transmission. Vet Rec. 2021 Apr;188(8):e247.

31. [31] Krafft E, Denolly S, Boson B, et al. Report of One-Year Prospective Surveillance of SARS-CoV-2 in Dogs and Cats in France with Various Exposure Risks: Confirmation of a Low Prevalence of Shedding, Detection and Complete Sequencing of an Alpha Variant in a Cat. Viruses. 2021 Sep 3;13(9).

32. [32] Sila T, Sunghan J, Laochareonsuk W, et al. Suspected cat-to-human transmission of SARS-CoV-2, Thailand, July-September 2021. Emerg Infect Dis. 2022 Jul;28(7):1485–1488.

33. [33] OIE. SARS-CoV-2 in animals - Situation Report 11. World Organisation for Animal Health 2022.

34. [34] Lineage Mutation tracker [Internet]. 2022 [cited 2022]. Available from: https://outbreak.info/situation-reports?pango=B.1.234.

35. [35] Khare S, Gurry C, Freitas L, et al. GISAID’s Role in Pandemic Response. China CDC Wkly. 2021 Dec 3;3(49):1049–1051.

